# A G-protein-coupled receptor mediates neuropeptide-induced oocyte maturation in the jellyfish *Clytia*

**DOI:** 10.1101/801225

**Authors:** Gonzalo Quiroga Artigas, Pascal Lapébie, Lucas Leclère, Philip Bauknecht, Julie Uveira, Sandra Chevalier, Gáspár Jékely, Tsuyoshi Momose, Evelyn Houliston

**Author notes:** GQA: The Whitney Laboratory for Marine Bioscience, University of Florida, St. Augustine, FL, USA. Corresponding author: E. Houliston.

## Abstract

The reproductive hormones that trigger oocyte meiotic maturation and release from the ovary vary greatly between animal species. Identification of receptors for these Maturation Inducing Hormones (MIHs), and understanding how they initiate the largely conserved maturation process, remain important challenges. In hydrozoan cnidarians including the jellyfish *Clytia hemisphaerica*, MIH comprises neuropeptides released from somatic cells of the gonad. We identified the receptor (MIHR) for these MIH neuropeptides in *Clytia* using cell culture-based “deorphanization” of candidate oocyte-expressed GPCRs. *MIHR* mutant jellyfish generated using CRISPR-Cas9 had severe defects in gamete development or in spawning both in males and females. Female gonads, or oocytes isolated from *MIHR* mutants, failed to respond to synthetic MIH. Treatment with the cAMP analogue 5’Br-cAMP to mimic cAMP rise at maturation onset rescued meiotic maturation and spawning. Injection of inhibitory antibodies to Gα_S_ into wild type oocytes phenocopied the *MIHR* mutants. These results provide the molecular links between MIH stimulation and meiosis initiation in hydrozoan oocytes. Molecular phylogeny grouped *Clytia* MIHR with a subset of bilaterian neuropeptide receptors including Neuropeptide Y, Gonadotropin Inhibitory Hormone, pyroglutamylated RFamide and Luqin, all upstream regulators of sexual reproduction. This identification and functional characterisation of a cnidarian peptide GPCR advances our understanding of oocyte maturation initiation and sheds light on the evolution of neuropeptide-hormone systems.

## Introduction

Oocyte meiotic maturation is an essential process for animal sexual reproduction. It transforms the tetraploid, fully-grown, ovarian oocyte into a haploid female gamete [1]. The core biochemical and cellular pathways operating within the oocyte during maturation are highly conserved between animals across the phylogenetic spectrum. Activation of the CyclinB-Cdk1 kinase complex assures meiotic progression from prophase I arrest into M phase, while parallel Mos-MAP kinase activation steers polar body formation as well as cytostatic arrest once maturation is complete [2,3]. In contrast the upstream physiological processes vary widely, as does the molecular nature of the maturation inducing hormones (MIHs) that act on the oocyte to trigger maturation [1,4]. This reflects clade-specific acquisition of endocrine tissues such as ovarian follicles, the corpus cardiacum/corpus allatum in insects, or the pituitary in vertebrates [5,6]. These tissues act downstream of other neuroendocrine sites such as the vertebrate hypothalamus to integrate environmental, behavioural and physiological information in order to achieve optimal conditions for gamete development and release [7]. The complex evolutionary history of hormonal reproductive regulation has made it challenging to unravel the crucial regulatory events operating within the oocyte at the onset of maturation.

G-protein coupled receptors (GPCRs), the largest superfamily of integral transmembrane receptors [8], are good candidates to serve as MIH receptors. These 7-transmembrane domain proteins activate a variety of cytoplasmic signalling pathways via Gα and/or Gβγ subunits that become released from receptor-associated heterotrimeric Gαβγ protein upon ligand binding [9–11]. Members of the four main Gα subunits classes, Gα_s_, Gα_i_ Gα_q_ and Gα_12/13_, associate variously with members of the vast GPCR family. In vertebrates, constitutively active GPCRs are coupled to Gα_s_, which stimulates adenylate cyclase to maintain high cytoplasmic cAMP concentrations in ovarian oocytes [12–14]. These high cAMP levels help hold the oocyte in an immature state, with the cell cycle arrested in meiotic prophase. The mechanisms by which MIHs override this arrest and initiate meiotic maturation vary between species and are not fully understood (see Discussion). In marked contrast, oocytes of many invertebrate species show a rise in cytoplasmic cAMP concentration upon MIH stimulation that is required for meiotic maturation [6]. In these species, GPCRs working through Gα_s_ are thus good candidates to trigger oocyte maturation, rather than to inhibit it as in vertebrates. The identification of such receptors could help to understand the origin of this diversity by providing an evolutionary perspective.

Here we identify the MIH receptor (MIHR) in the hydrozoan jellyfish *Clytia hemisphaerica* as a GPCR likely working through Gα_s_, and determine its *in vivo* function. In *Clytia* and other hydrozoan species, MIH consists of PRPamide and related tetrapeptides that are released upon light stimulation from opsin-expressing gonad ectoderm cells, and act at the plasma membrane [15,16]. Neuropeptides such as these are of major importance in cnidarian biology, acting both as “neuroendocrine” mediators of physiological transitions such as metamorphosis, as well as in fast neuromuscular transmission regulating swimming and feeding mediated by ligand-gated ion channels [reviewed by 17,18]. We uncovered an oocyte-expressed Class A GPCR in *Clytia* (MIHR) activated by MIH peptides, which is to our knowledge the first characterized cnidarian neuropeptide GPCR [19,20]. Antibody inhibition experiments further provided evidence that Gα_s_ links MIHR activation to cAMP production to trigger maturation [21,22]. CRISPR-Cas9 mediated mutation of the *MIHR* gene revealed an essential *in vivo* function of *MIHR* in initiating oocyte maturation. Phylogenetic analysis of the MIHR sequence identified an evolutionary link to a subset of bilaterian neuropeptide-hormone GPCR families, including several upstream regulators of reproduction. These results allow us to propose a new scenario for the evolution of hormonal signalling pathways regulating oocyte maturation.

## Results

### Selection of candidate MIH GPCRs

Amidated neuropeptides like MIH commonly signal through GPCRs, although other receptor types can also be used [23]. As the first step to select candidate MIH receptors, we compiled a comprehensive catalogue of *Clytia* GPCRs from a *Clytia* reference transcriptome covering all life-cycle stages. Our bioinformatics pipelines first retrieved all sequences predicted to code for 7 transmembrane domain (7TM domain) proteins and bearing GPCR-related Pfam tags. An initial list of 761 sequences was then assigned by Pfam to the three main GPCR classes: A (rhodopsin-like), B (secretin-like) and C (metabotropic glutamate-like)[8,24] or to an “other” category (which included for instance sweet-taste receptors and cAMP-like receptors). The final dataset of 536 class-sorted putative *Clytia* GPCRs obtained after removal of duplicates (File S1) may be a slight overestimate due to some incorrectly identified or incomplete sequences. We focussed on the 377 class-A GPCRs, since most neuropeptide GPCRs belong to this class [8, 25].

An important criterion for the selection of MIH-receptor candidates from the class-A GPCR list was enrichment in oocytes (Fig.1A). We mapped Illumina HiSeq mRNA reads previously obtained from *Clytia* gonad tissues [16] and life cycle stage [26] against all putative GPCR sequences. Profile clustering revealed three groups of sequences with oocyte-enriched expression (Fig. S1). We further narrowed down the number of potential MIHRs from these expression groups to 96 sequences, taking into account also Pfam indicators and sequence similarity with a set of bilaterian GPCRs [25; see Figure 6 for clustering of a subset of sequences closest to the *Clytia* MIHR]. Finally we compiled a short-list of 16 candidates for functional testing, using high expression level in oocytes as the final selection criterion (Fig.1A, File S2).

**Figure 1.**
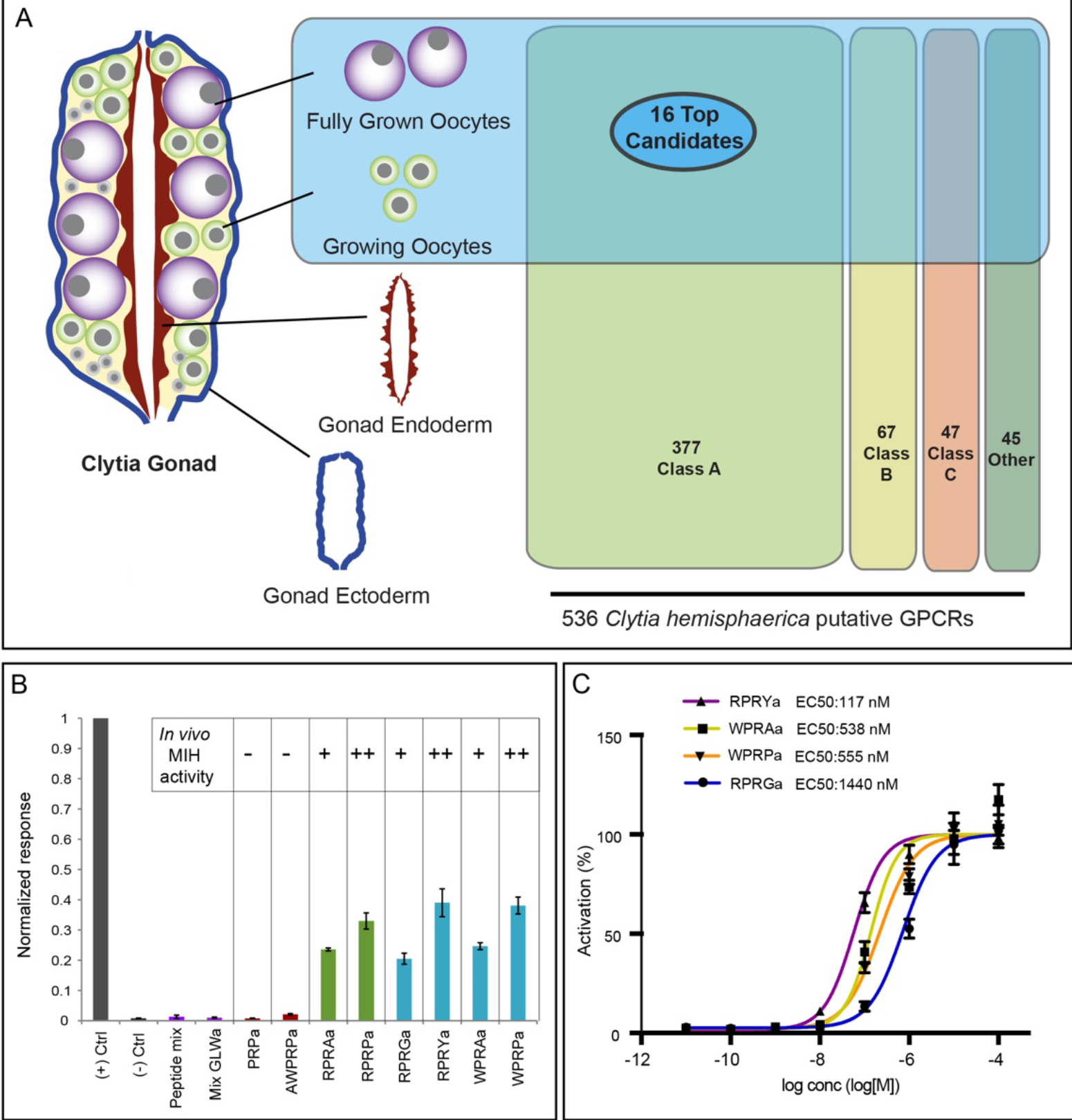
Identification of the *Clytia* MIH receptor (MIHR) **A)** Diagram of a *Clytia* gonad showing the different tissues used for RNA-seq and GPCR expression comparisons. 536 putative GPCRs identified in *Clytia* mixed-stages transcriptome were assigned to the three main GPCR classes (A, B, C) or “other” based on Pfam signatures. 16 top candidate MIH GPCRs were selected based on oocyte enrichment, relatedness to known bilaterian class-A neuropeptide GPCRs and Pfam information. **B)** Luminescence response of CHO-K1 cells expressing the putative *Clytia* MIH GPCR treated with neuropeptide mixes lacking MIH activity (purple bars), MIH tetrapeptides identified from *Clytia hemisphaerica* (blue bars) or *Cladonema radiatum* (green bars), or related penta- and tripeptides previously shown to be ineffective in triggering oocyte maturation *in vivo* (red bars) [15]. Empty pcDNA3.1 vector was used as negative control, and *Platynereis* FLamide and its receptor as a positive control (grey bars). Peptide concentrations all 1μM. Absolute units of luminescence were normalized using the positive control; data are shown as mean ± standard error of the mean (n= 3). MIH tetrapeptides were selectively able to activate the *Clytia* GPCR, the responses closely matching the *in vivo* MIH activity of each peptide tested on *Clytia* oocytes as indicated [summarised results from 15]. **C)** Dose–response curves of *Clytia* MIHR challenged with four variant *Clytia* MIH tetrapeptides. One of 3 independent experiments with equivalent results is shown. Luminescence values were normalized relative to the maximum of fitted dose–response curves, and are shown as mean ± standard error of the mean (n= 3). Half maximal effective concentration (EC50) values were calculated as means of 3 independent experiments.

### Identification of the *Clytia* MIH receptor by GPCR deorphanization assay

We used a cell culture-based GPCR deorphanization assay to identify the MIH receptor from our candidate shortlist. cDNAs for each candidate were transfected into CHO-K1 cells along with an aequorin-GFP luminescence reporter that measures Ca2+ mobilization downstream of a promiscuous Gα protein [27,28]. We first screened the 16 candidate *Clytia* GPCRs against a mixture of 33 synthetic amidated peptides (including MIHs) predicted to be generated from previously identified putative *Clytia* neuropeptide precursors [15] and one additional one identified from transcriptome data (File S4). Given our imperfect knowledge of pro-peptide processing in cnidarians, some of the synthetic peptides may not correspond to endogenous peptides [29,30]. Only one of the 16 GPCRs was activated by this peptide mixture. This receptor responded well to all 4 synthetic *Clytia* MIH tetrapeptides, as well as to related *Cladonema* MIH peptides that can also activate *Clytia* oocyte maturation, but not to other *Clytia* peptide mixtures or poorly-active MIH penta-/tripeptides. We termed this *Clytia* GPCR the Maturation Inducing Hormone Receptor (MIHR). The activity of individual peptides at 1 μM to stimulate the MIHR closely matched their *in vivo* ability to induce oocyte maturation [15; Fig.1B].

The *Clytia* MIH amidated peptides are produced from two precursor genes *Che-pp4* (generating multiple copies of WPRAa, WPRYa and WPRPa) and *Che-pp1* (multiple copies of WPRPa and RPRGa) [15; File S4]. Their predicted structural similarity suggests they bind the same site on the receptor with different affinities. Dose-response curves for each of the 4 *Clytia* MIH neuropeptides, generated from three independent experiments, showed half-maximal effective concentration (EC50) values in the high-nanomolar or low-micromolar range for all 4 MIHs, with RPRYamide showing the highest activity and RPRGamide the lowest (Fig. 1C). To our knowledge, *Clytia* MIH and MIHR are the first neuropeptide ligand-GPCR pair demonstrated in a cnidarian.

### Slow growth, gametogenesis disruption and spawning failure in *MIHR* mutant jellyfish

To determine the function of MIHR *in vivo* we generated *Clytia MIHR* knockout (KO) polyp colonies using CRISPR/Cas9 mutagenesis (see Methods). This gene-editing technique is very effective in *Clytia*, allowing extensive bi-allelic mutation of target genes already in F0 polyp colonies [16, 63]. Three guide RNAs designed to target the 3^rd^ transmembrane domain of the MIHR protein were tested by genotyping at the planula stage, and the most effective one used to generate polyp colonies. After genotyping around the target site, we selected and propagated for phenotypic analysis six colonies showing no detectable wild type sequence and carrying mainly frame-shift mutations (Table 1). All six mutant polyp colonies expanded slowly compared to wild type colonies. They displayed variable morphologies, with two of them rambling pronouncedly away from the glass substrate (Fig. 2A; Table 1), but all produced active gonozooids that budded baby medusae. Growth of these mutant baby medusae was slower for the six mutants than for wild types, but they were able to reach the full adult size of about 1cm in diameter. The adult jellyfish all swam less vigorously than wild types. The slow growth of mutant polyps and medusae was not due to any obvious feeding problems: both forms could capture *Artemia* nauplii without difficulty.

**Figure 2.**
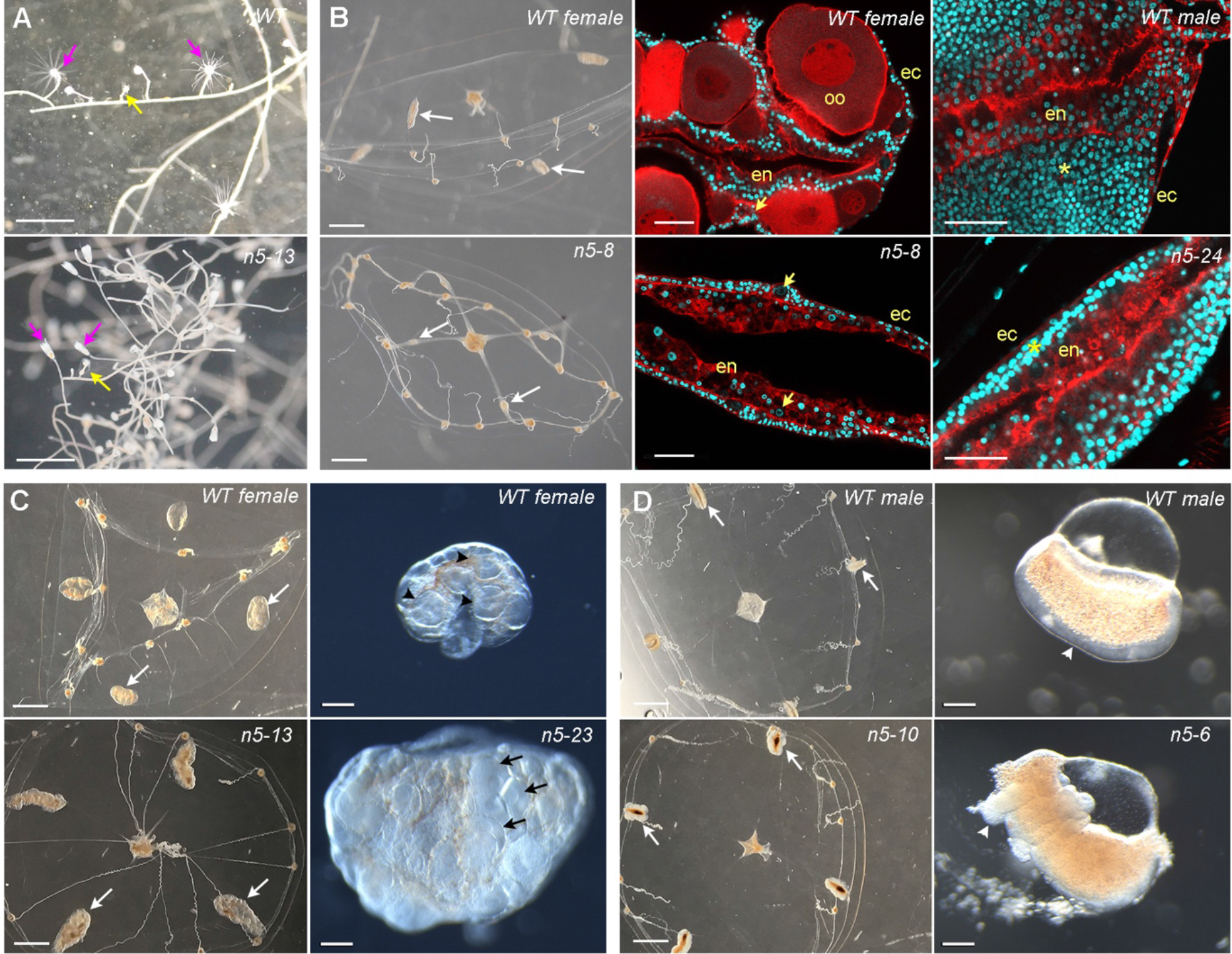
Phenotypes of *Clytia MIHR* mutants. Light and confocal microscope images of *MIHR* mutant F0 polyp colonies and jellyfish. **A)** Morphology of a wild type (WT) colony (Z11, top) and a *MIHR* mutant colony (n5-13, bottom). All *MIHR* mutant colonies contained gastrozooid and gonozooid polyps (pink and yellow arrows respectively), however the connecting stolons in some mutant colonies (see Table 1) were convoluted and frequently detached from the glass substrate, while stolons of wild type colonies were straight and adhered tightly. **B)** Fully grown mutant jellyfish n5-8 and n5-24 (bottom row) compared to wild type (top row) showed poorly developed gonads (white arrows) and sluggish behaviour but are otherwise indistinguishable. Centre (females) and right (males) panels show confocal microscope images through th gonads of adult medusae; Nuclei are stained with Hoechst 33342 (cyan) and F-actin with Phalloidin (Red). In the n5 8 female gonad, small oocytes (arrows) can be detected between the endodermal (en) and ectodermal (ec) layers, but no large growing oocytes (oo) are present compared to the wild type female gonad. In the n5-24 male gonad, the spermatogenic zone (asterisk) between endoderm and ectoderm is much thinner than in the wild type male gonad. **C)** Comparison of mutant n5-23 and n5-13 (bottom row) female medusae gonads (white arrows) swollen by an accumulation of large oocytes to wild type (Z11) female medusae (top row). Right panels show gonads dissected from 3-week old n5-23 medusae 10 hours after a light cue that induced spawning in the wild type but not the mutant gonad. **D)** Fully grown mutant n5-10 and n5-6 male medusa (bottom row) compared to wild type (Z13) male medusae (top row) have deformed gonads (white arrows). Right panels show isolated gonads, illustrating the thickened and irregular spermatogenic layer (arrowheads) in the mutant. Scale bars 1mm for light microscope jellyfish and polyp images; 200μm for isolated gonad images; 50μm for confocal images.

**Table 1.** Characteristics of *MIHR* mutant colonies

| **Colony name** | **Predominant mutations** | **Medusa sex** | **Polyp colony\*** | **Medusa gonads** |
| --- | --- | --- | --- | --- |
| n5-23 | -4nt , +74nt | F | normal | Enlarged and deformed |
| n5-8 | -7nt | F | rambling | Poorly developed |
| n5-13 | -4nt , -3nt | F | rambling | Enlarged and deformed |
| n5-24 | -34nt , +59nt | M | normal | Poorly developed |
| n5-10 | -4nt | M | normal | Enlarged and deformed |
| n5-6 | -4nt , -3nt | M | normal | Enlarged and deformed |

\*see Fig. 2A

Of the six *MIHR* mutant colonies, three produced male jellyfish and three females. In addition to the shared defects in growth and swimming behaviour, they showed one of two distinct phenotypes affecting gonad development. For one female and one male *MIHR* mutant (n5-8 and n5-24 respectively), the gonads of the adult jellyfish developed poorly, failing to accumulate gametes as occurs during wild type adult growth (Fig. 2B). Examination of the gonads by confocal microscopy revealed the presence of small oocytes for n5-8 and a thin spermatogenic zone for n5-24. This suggests that mutation of the *MIHR* gene can compromise gametogenesis in male and female jellyfish. One possible explanation is that gamete development can stall completely when overall growth of the jellyfish is poor. In marked contrast, jellyfish from the four other *MIHR* mutant polyp colonies underwent gametogenesis to produce large numbers of fully-grown oocytes (n5-13, n5-23; Fig. 2C) or spermatozoids (n5-6, n5-10; Fig. 2D). In the female medusae, the gonads were grossly inflated by accumulation of fully-grown immature oocytes, due to failure in light-induced oocyte maturation and subsequent release of the unfertilised egg (see below). In males, the gonads were markedly deformed and irregular. Following dark-light transitions that induced spawning in wild type males, some local sperm release was observed from rupture sites of the gonad epithelium, but the sperm remained concentrated at the gonad surface and failed to disperse.

Overall this initial phenotype analysis of six independent *MIHR* mutant F0 colonies indicated that MIHR has non-essential functions in medusa growth and in regulating medusa swimming. These could relate to expression of MIH and MIHR at other sites in the jellyfish, notably the manubrium (mouth/stomach structure) and tentacles (see below). In addition, two distinct gonad defects were observed: failure of gamete development or over-accumulation of mature gametes. These had no obvious relationship to particular genotypes of the corresponding F0 polyp colonies (Table 1) and so may be the consequence of *MIHR* mutation in different genetic or epigenetic backgrounds. In any case, the accumulation of immature, fully grown oocytes in medusae from two independent *MIHR* mutants strongly supported a key role for this GPCR in oocyte maturation, as characterized further below.

### *Clytia* MIHR is expressed in oocytes and also in tentacle cells

The defects observed in behaviour and growth as well as in gametogenesis in MIHR mutant jellyfish suggested that this receptor may have roles both in the gonads and other sites. *In situ* hybridization detection of *MIHR* mRNA in adult jellyfish (medusae) and in isolated gonads (Fig. 3A, B) supported this idea. Strong expression was expressed in developing oocytes and in male gametes, consistent with the ability of MIH to induce male spawning [15]. Expression was also detected in clusters of small somatic cells located within each tentacle bulb. Individual cells from these clusters extended in a line along each tentacle. The location of these cells suggests that they belong to a neural subtype potentially involved in regulating tentacle contraction. The MIH receptor in these tentacle cells could potentially bind MIH neuropeptides produced by neural/neuroendocrine cells of the endodermal gastrovascular system, detectable using an antibody recognising the PRPamide peptides produced from both Che-pp1 and Che-pp4 [anti-PRPa; 15; Fig. 3C]. These two precursors are expressed respectively exclusively in the manubrium and tentacle, as well as being co-expressed in scattered cells of the gonad [15]. As well as the cells in the gonad ectoderm that mediate spawning, the anti-PRPa antibody decorates cells in the tentacle especially at its junction with the circular gastrovascular canal, and in the endoderm of the manubrium feeding/digestive organ [15,16] (Fig. 3C). Comparison of *in situ* hybridization patterns for *MIHR* and *MIH* showed that the ligand and receptor-expressing cells in the tentacles have distinct distributions: the single file of MIHR cells lies on the rounded oral side [see 31], whereas the MIH cells form two flanking lines (Fig. 3B). The position and morphology of these cells suggest that they correspond to two sub-populations of neural cells positioned between the tentacle endoderm and ectoderm in hydrozoans [31,32]. No *MIHR*-expressing cells were detected in the manubrium, only non-specific staining of the manubrium floor. Our inability to detect MIHR expression close to these manubrium PP1-expressing cells may be due to the limitations of the *in situ* hybridization technique, with low expression in the manubrium and/or other tissues not sufficient to provide a clear signal. Another possibility is that a second receptor is expressed in the manubrium and more sensitive to RPRGa/WPRPa peptides. Alternatively, the *MIHR*-expressing cells at the base of the tentacles might respond to endocrine MIH tetrapeptides circulating in the gastrovascular system, for example secreted in response to nutritional status. Regulation by these cells of tentacle contractions, and directly or indirectly of swimming behaviour, could account for the sluggish movement and poor growth of *MIHR* mutant medusae. Specific assays should be devised in the future to examine this hypothesis in detail.

**Figure 3.**
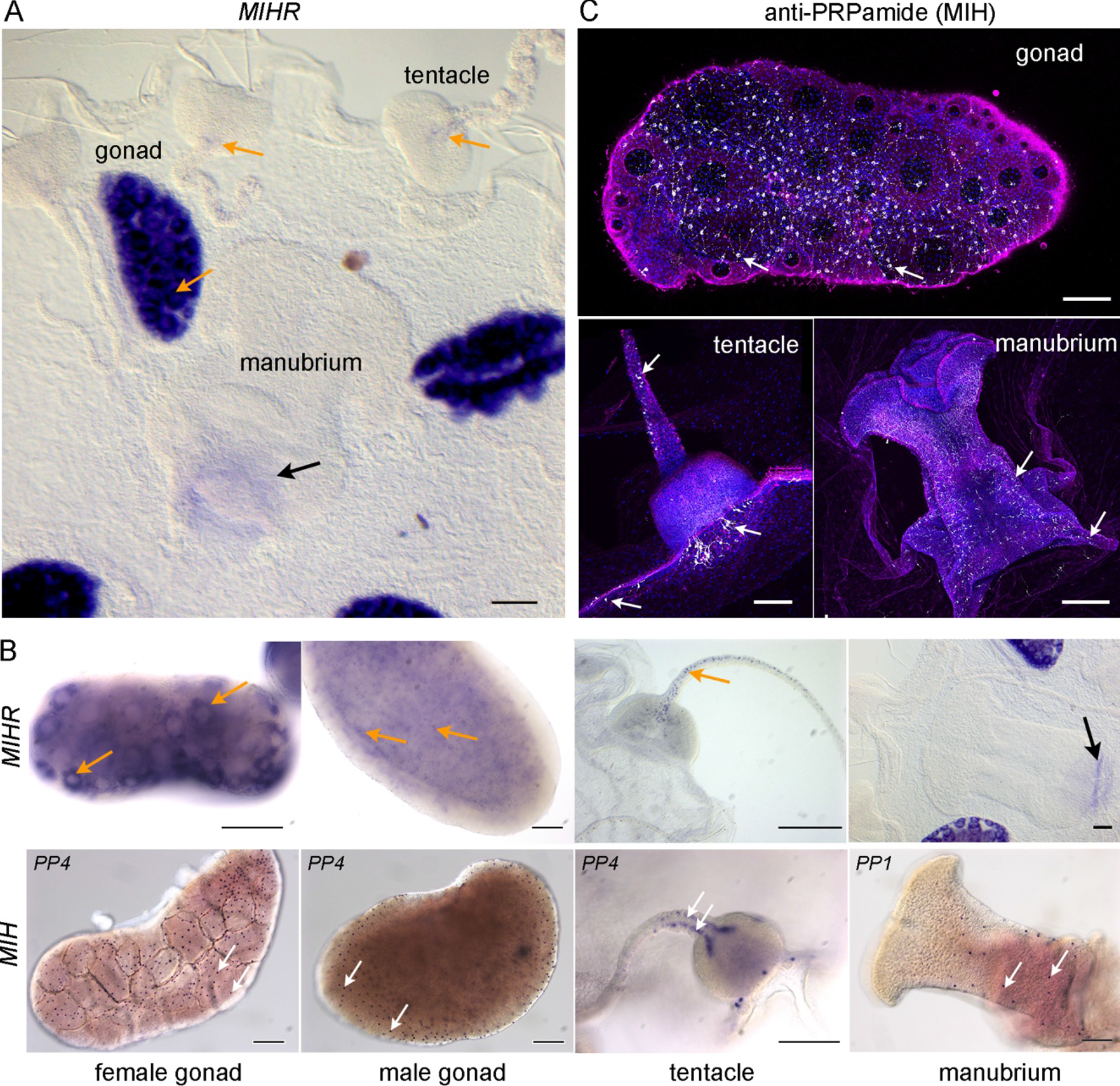
Sites of *MIHR* expression in the *Clytia* medusa. **A)** *In situ* hybridization detection of *MIHR* mRNA in a *Clytia* female medusa. Strong purple *MIHR* signal (orange arrows) was detected in oocytes within the gonads, as well as in scattered cells in tentacles. **B)** Comparison of the distribution of the MIH receptor and ligand expressing cells in different medusa structures as labelled, detected by *in situ* hybridization using probes to *MIHR* (top row) and to the MIH peptide precursors *PP1* and/or *PP4* as indicated (bottom row). Orange arrows point oocytes, developing spermatozoa and tentacle MIHR cells, and white arrows indicate MIH cells. The focal plane in the male gonad image is through the centre to illustrate the position of the MIH cells in the ectodermal layer. Weak staining at the base of the manubrium (black arrow) in A and B is frequently observed with probes for many genes and is probably due to a specific trapping of the colour reagent. Scale bars: 100 μm. **C)** Confocal images of the three main sites of MIH-expressing cells (white arrows) in medusae, visualized using anti-PRPamide antibody (MIH: white), anti-tyrosinated tubulin (magenta) and Hoechst staining of nuclei (blue). Summed Z stacks are shown in all cases except for the gonad tubulin and DNA staining, where a single plane was selected through the centre of the gonad. All scale bars 100μm.

### *Clytia* MIHR is the essential oocyte receptor for oocyte maturation

We used jellyfish from the two *MIHR* female mutant colonies showing the swollen-gonad phenotype (n5-13 and n5-23) to characterize the role of this GPCR in oocyte maturation. For n5-23 the predominant genotypes were a 4 base deletion which introduces a premature STOP codon, and a 55 base insertion, but a small proportion of wild type cells was also detected. Colony n5-13 contained a majority 4 base deletion but also a high proportion of a 3 base deletion. We performed a series of oocyte maturation and spawning assays, comparing the responses of wild type and *MIHR* KO isolated gonads and oocytes to different stimuli (Figure 4). Isolated wild type gonads underwent oocyte maturation and spawning in response to light stimulation or treatment with the synthetic MIH peptide WPRPamide (100nM) as previously shown [15], but *MIHR* mutant gonads did not respond (Fig. 4A, B). In contrast, treatment with 4mM 5Br-cAMP, a cell-permeable analogue of cAMP which induces hydrozoan oocyte maturation by mimicking the early cytoplasmic cAMP rise [3,21], rescued the phenotype of *MIHR* mutant gonads, efficiently triggering oocyte maturation and spawning. 5-Br-cAMP treatments but not MIH peptides also promoted maturation of isolated fully grown *MIHR* mutant oocytes (Fig. 4C). Br-cAMP-matured *MIHR* mutant oocytes could be fertilized to develop into planula larvae, although had lower development and metamorphosis rates than wild type oocytes.

**Figure 4.**
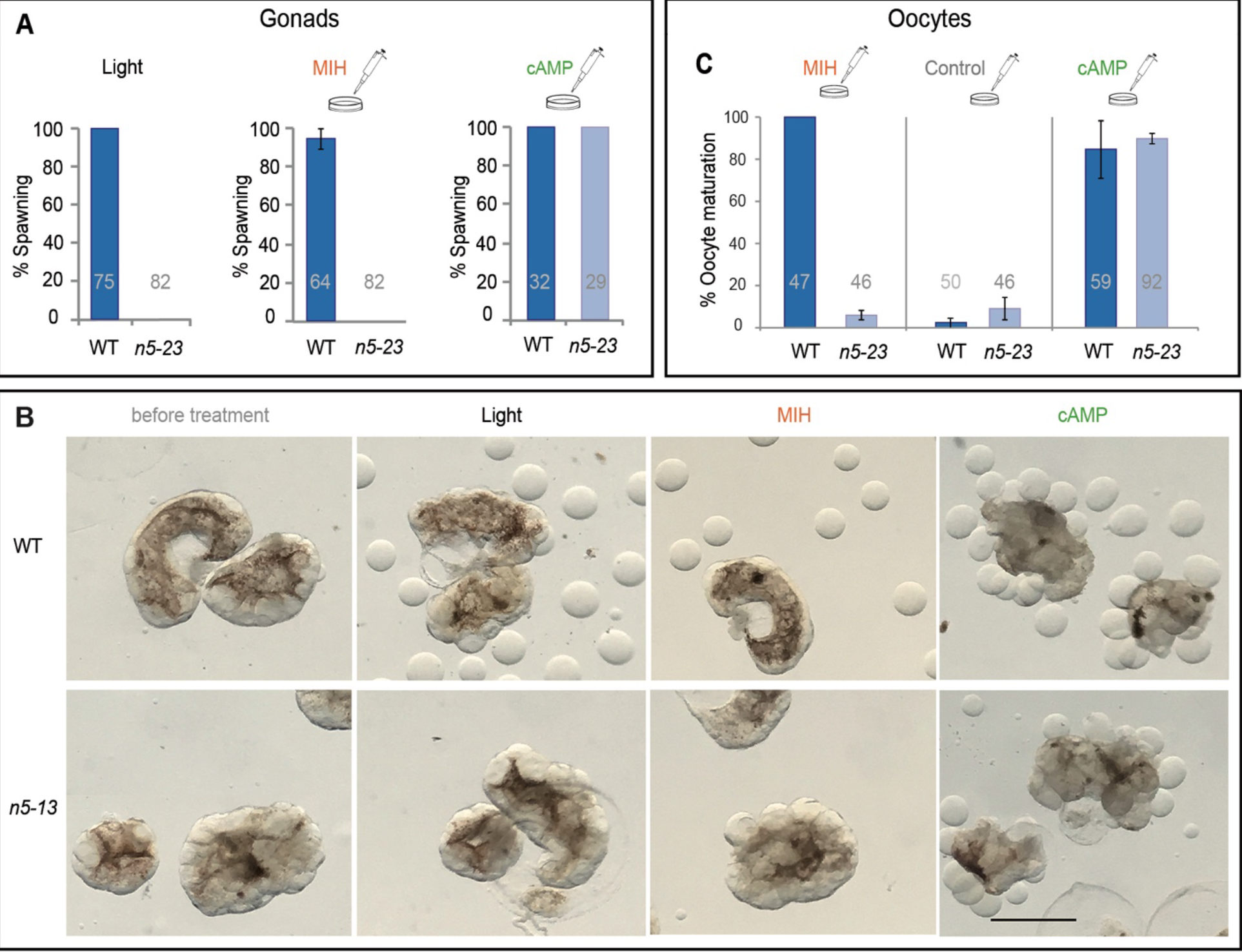
Oocyte maturation failure in *MIHR* mutant medusae. Oocyte maturation assays performed using isolated gonads (A, B) or isolated oocytes (C) from *MIHR* mutant jellyfish compared to wild type (WT). In A and C, bar heights represent mean percentages of 3 independent experiments and error bars show standard deviations. Total gonad or oocyte numbers for each treatment are indicated in grey. **A)** Spawning response of isolated gonads from WT and n5-23 female medusae. Three treatments were compared as indicated above each panel: Light: light stimulation after incubation in the dark; MIH: treatment of light-maintained gonads with 100nM WPRPamide; cAMP: treatment of light-maintained gonads with 4mM 5-Br-cAMP. No oocyte maturation or spawning were observed in *MIHR* KO gonads upon light stimulation or MIH treatment, while 5-Br-cAMP treatment provoked oocyte maturation and spawning. The Fisher exact test showed significant differences (F= 0) between wild type and mutant responses to light and MIH, but not for the cAMP treatment (F=1). **B)** Light microscope images illustrating gonads from an equivalent experiment performed with n5-13 female *MIHR* mutant medusae 120 minutes after the indicated treatments. Scale bar 500μm. **C)** Response of fully grown oocytes isolated from WT and n5-23 *MIHR* mutant gonads to MIH and 5-Br-cAMP treatments as in A. Both treatments triggered maturation of WT oocytes, visible after 20-30 minutes as germinal vesicle breakdown (GVBD), but only 5-Br-cAMP induced maturation of *MIHR* KO oocytes. Control experiments using the 5-Br-cAMP solvent (distilled water) showed a low level of spontaneous maturation in both cases. Fisher exact test did not show significant differences between WT and mutant oocytes in the control (F= 0.101) or cAMP treated (F= 0.216) groups, but did so in the MIH assays (F= 0).

This demonstration of a cAMP-reversible maturation initiation failure in mutant female jellyfish confirmed that the *Clytia* MIHR has an essential *in vivo* function as the oocyte receptor for MIH.

### MIH-induced oocyte maturation blocked by an inhibitory Gα_s_ antibody

GPCR activation can lead to a cytoplasmic cAMP concentration rise in the responding cell as a result of adenylate cyclase stimulation via S type Gα subunits of heterotrimeric G proteins. We tested the role of Gα_s_ in *Clytia* oocyte maturation by injecting isolated oocytes with a specific inhibitory antibody previously shown to cause meiotic maturation of mouse, *Xenopus* and zebrafish oocytes [12,33,34]. All oocytes were tested for maturation competence by incubation at the end of the experiment in 5’-Br-cAMP. Any that failed to mature when treated with cAMP were discounted in the analyses. Oocytes injected with anti-Gα_s_ responded less efficiently than oocytes injected with PBS or a control anti-GST antibody when treated with synthetic MIH at low and then high doses (10nM then 100nM WPRPamide; Kinetics observed in one of three equivalent experiments shown in Fig. 5A). These results strongly suggest that MIHR acts mainly through Gα_s_.

**Figure 5.**
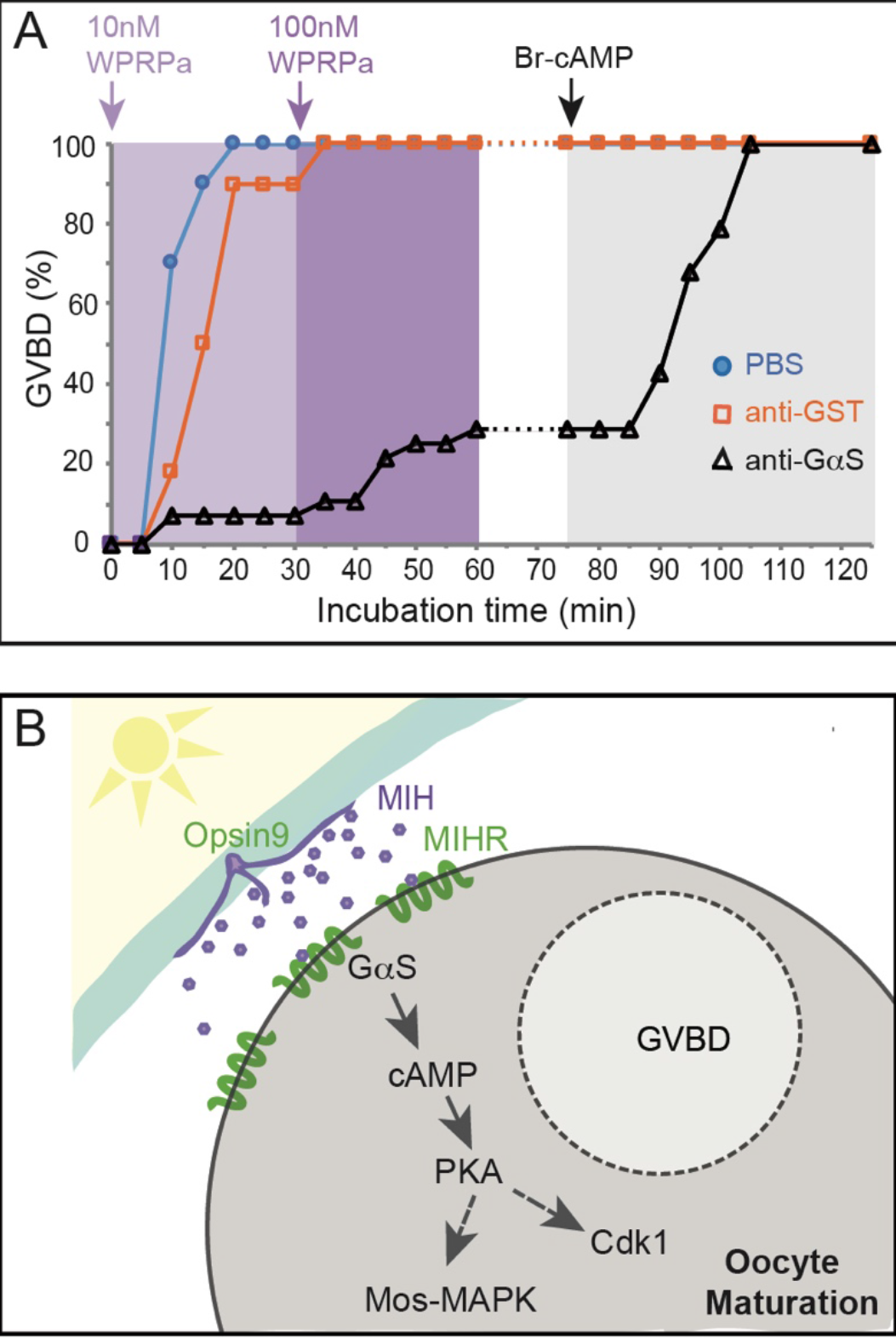
MIH-induced oocyte maturation blocked by an inhibitory Gα_s_ antibody. Involvement of Gα_s_ in *Clytia* oocyte maturation. **A)** Results of an antibody inhibition experiment. Maturation response (scored as % GVBD over time) of isolated oocytes injected with antibodies or buffer, then challenged first with a low dose of MIH (10nM WPRPamide), then with a higher dose (100nM WPRPamide) and finally with 5’-Br-cAMP to verify maturation competence, as indicated by the coloured arrows and background shading. Oocytes injected with PBS (blue circles) or a control anti-GST antibody (orange squares) responded efficiently to MIH, whereas very few oocytes injected with anti-Gα_s_ (black triangles) underwent GVBD after treatment with the low dose of MIH and only 29% following the high dose. The number of oocytes per group in this experiment was 30, 28 and 30 respectively. All of them had undergone GVBD by the end of the experiment. Times from the start of the first incubation are shown on the x axis. Equivalent results were obtained in two other experiments. **B)** Scheme illustrating the proposed cascade initiating *Clytia* oocyte maturation initiation. Following light stimulation after a dark period, Opsin9 mediates release of MIH neuropeptides from specialised cells (purple) of the gonad ectoderm (cyan). Activation of the MIHR (green) at the oocyte surface releases Gα_S_ to promote an increase in cytoplasmic cAMP, activating PKA. Unknown PKA substrates likely trigger in parallel Cdk1 activation and thus GVBD, and Mos1 synthesis to initiate the MAPK cascade.

Based on these findings we can now propose a model for maturation initiation in *Clytia* oocytes through MIH, MIHR, Gα_s_ and cAMP (Fig. 5B). A light cue triggers MIH release from neuroendocrine-type cells in the gonad ectoderm via an essential opsin protein [16]. The MIH peptides act on MIHR at the oocyte surface to promote Gα_s_ activation, stimulating adenylate cyclase and thus promoting an increase in cytoplasmic cAMP concentration, essential for the transition into meiotic M phase.

### *Clytia* MIHR is related to a bilaterian superfamily of neuropeptide hormone receptors

Much progress has been made in understanding the relationships between GPCR-neuropeptide families between protostomes and deuterostomes [20,25,28,35,36], and more recently with those in acoels and cnidarians [37]. We performed sequence-similarity based clustering to explore the relationship of the *Clytia* MIHR sequence with known GPCR families in Bilateria using established datasets from human and the annelid *Platynereis* [25,28 Fig. 6A]. The *Clytia* MIHR fell within the majority group of cnidarian GPCRs that cluster with a superfamily of peptide hormone receptors including human neuropeptide Y, neuropeptide FF, Tachykinin, Orexin/allastropin, Elevenin, and EFLGa/Thyrotropin releasing hormone, as well as Luqin from *Platynereis* (Fig. 6B). Adding to this analysis sequences recovered from the anthozoan cnidarian *Nematostella* highlighted the extensive independent expansion of GPCR-A sequence since cnidarian and bilaterian neuropeptide systems split [37]. This analysis demonstrated that the MIHR group does not contain any of the GPCRs directly involved in controlling oocyte maturation in vertebrates, including the constitutively active GPR3, gonadotropin releasing hormone (GnRH) receptor or Luteinizing/Follicle-stimulating hormone (LH/FSH) receptor (see Introduction and Figure 7). A GPCR from *Nematostella* annotated as a GnRH receptor [38] belongs as sister to distinct ‘superfamily’ including the bilaterian GnRH, vasotocin, CCAP corazonin and achatin receptor families.

**Figure 6.**
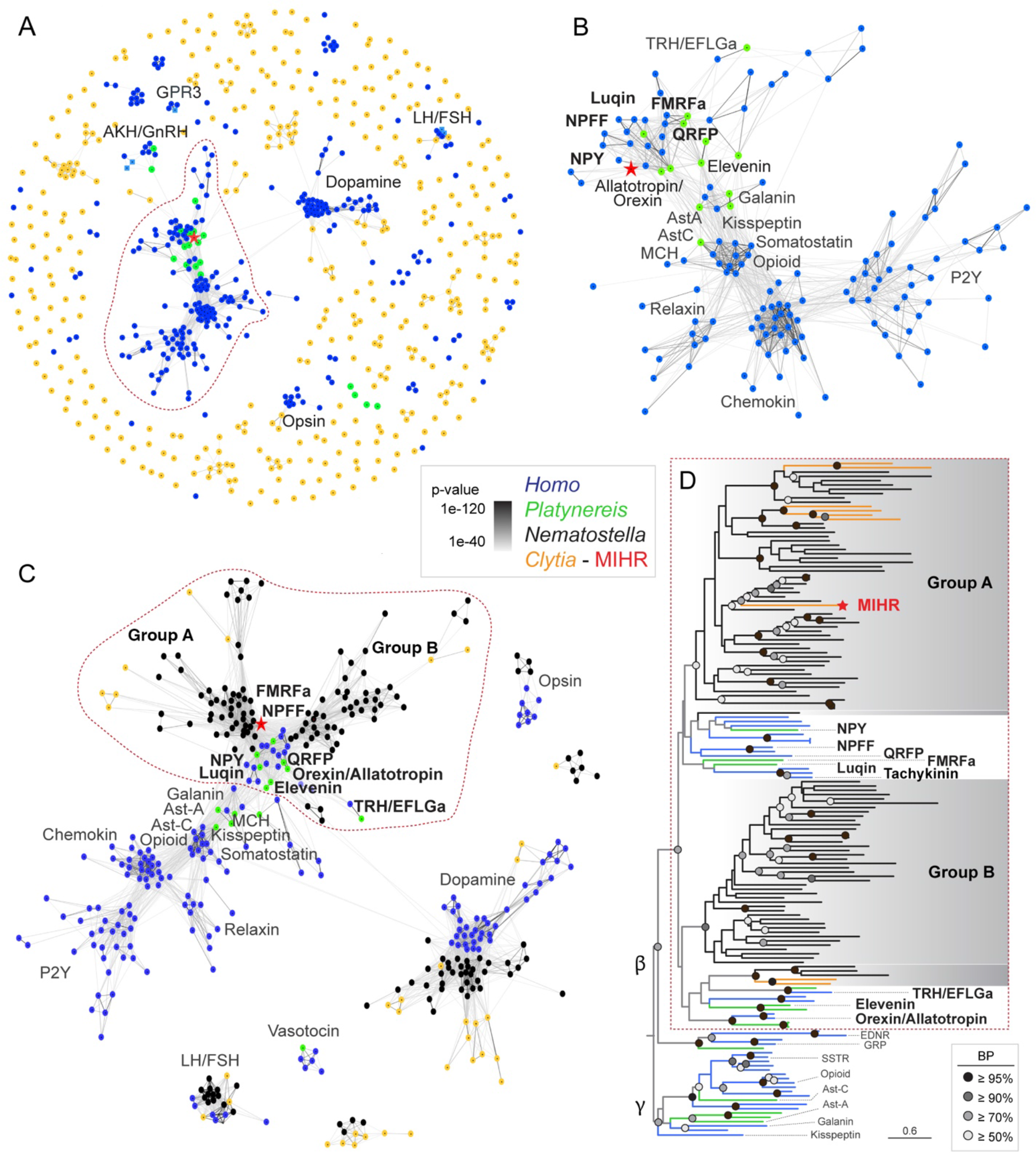
Relationship of *Clytia* MIHR to bilaterian neuropeptide hormone GPCRs. (A) Sequence-similarity-based clustering using Clans2 of all identified class-A GPCRs from *Clytia*, human (olfactory receptors excluded) and *Platynereis* deorphanized GPCRs [28] BLASTP p-value < 1e-40. (B) Cluster map of the largest cluster (circled in red in A) keeping only sequences that show at least 2 connections with the central cluster. BLASTP p-value < 1e-40. (C) More stringent cluster map (p-value < 1e-50) of the same sequences as in (A) plus all *Nematostella* GPCR-A sequences. Only clusters containing at least 5 sequences from at least 2 species were kept. All connections with p-value < 1e-40 are shown. (D) Maximum likelihood analysis of the sequences contained inside the dashed area shown in (C) using RaxML (PROTGAMMAGTR) with 500 Bootstrap replicates (BR). Tree file in Fig. S2. AstA, Galanin, AstC and Kisspeptin receptors were included as outgroup. AKH, adipokinetic hormone; AstC, allatostatin-C; AstA, allatostatin-A; GnRH, gonadotropin releasing hormone; LH/FSH, Luteinizing/Follicle-stimulating hormone receptor; GPR3, G protein-coupled receptor 3; MCH, melanin-concentrating hormone; NPY, neuropeptide Y; NPFF, neuropeptide FF; PRLH, prolactin releasing hormone; P2Y, purinoceptor; QRFP, pyroglutamylated RFamide peptide; TRH, thyrotropin releasing hormone. Colour code: *Homo sapiens*: blue, *Platynereis dumerilli:* green, *Nematostella vectensis*: black, *Clytia*: orange. Red star: *Clytia* MIHR.

**Figure 7.**
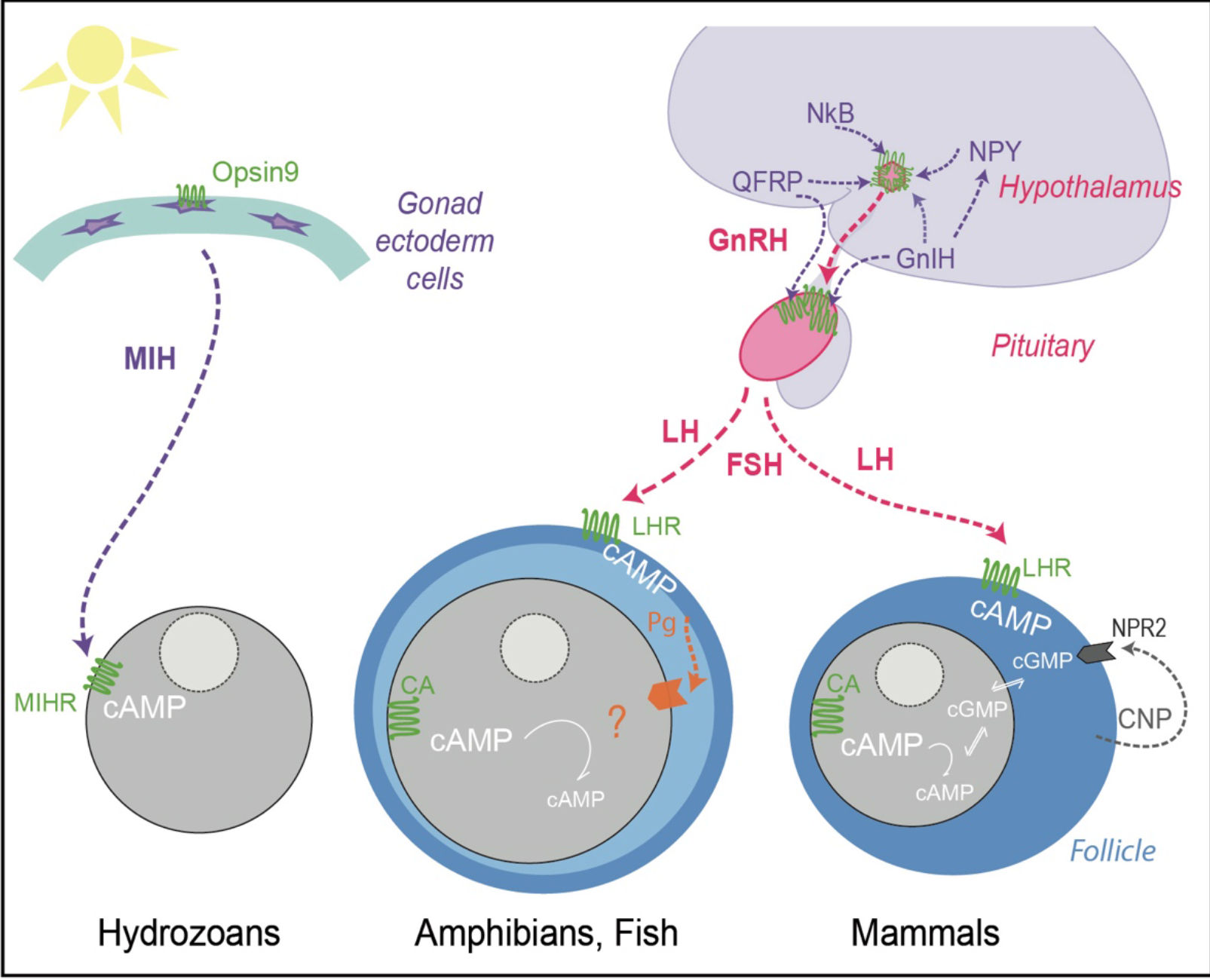
Schematic comparison of GPCR regulation of *Clytia* and vertebrate oocyte maturation. Simplified view of the tissues, hormones and receptors involved in regulating oocyte maturation in *Clytia*, and in fish/amphibians and mammals. For simplicity we have not included protostome or echinoderm models. The principle peptide hormones of the reproductive hypothalamus-pituitary-gonadal axis (GnRH and LH/FSH) are in pink, and those for which the receptors group phylogenetically with *Clytia* MIHR in “Group A” (Fig. 6) in purple. Peptide hormones: *Clytia* MIH; Neuropeptide Y (NPY); Gonadotropin Inhibitory Hormone (GnIH); Gonadotropin Releasing Hormone (GnRH); Luteinizing Hormone (LH); glutamine RF7amide peptide (QRFP); Neurokinin B (NkB); C-type natriuretic peptide (CNP). All their receptors, except the guanylyl cyclase natriuretic peptide receptor 2 (NPR2) activated by CNP, are GPCRs (green). Constitutively active (CA) GPCRs in vertebrate oocytes maintain cytoplasmic cAMP levels high prior to maturation. Several types of oocytes receptor (orange) may respond to steroid hormones (Pg) in different species of amphibians and fish, but the relative importance of multiple downstream signalling pathways remains to be clarified [1,43,73,74]. See text for discussion.

Maximum Likelihood phylogenetic analyses of the large GPCR cluster containing MIHR confirmed phylogenetic support for a GPCR ‘superfamily’ associating two distinct cnidarian groups (A and B in Fig. 6C, D) with a set of bilaterian neuropeptide hormone receptors [37]. Cnidarian GPCR group A contained *Clytia* MIHR, six other *Clytia* GPCRs and a large number of *Nematostella* receptors. It showed weakly-supported association with a set of bilaterian peptide hormone receptor families including those for neuropeptide Y/neuropeptide F, GnIH/neuropeptide FF, RYa/Luqin, Tachykinin and pyroglutamylated RFamide peptide (QRFP) [25,28,37]. Group B included two *Clytia* GPCRs and associated with further bilaterian neurohormonal receptor families including Elevenin, EFLGa/TRH and Orexin/Allatotropin receptors (Fig.6C,D). The low support values for many deep branch relationships within this superfamily makes it difficult to determine definitive evolutionary relationships between the neuropeptide hormone receptors and cnidarian group A and B GPCRs. We can nevertheless conclude that *Clytia* MIHR along with eight other *Clytia* GPCRs and a very large group of *Nematostella* GPCRs have a common ancestor with a subset of bilaterian peptide hormone receptor families that includes many involved in regulating sexual reproduction and feeding (see Discussion).

## Discussion

GPCR deorphanization followed by targeted gene mutation allowed us to identify *Clytia* MIHR as the neuropeptide receptor responsible for initiating oocyte meiotic maturation. *MIHR* female mutant medusae accumulated fully-grown oocytes that failed to mature unless treated with cAMP analogues to bypass the receptor. We further provided evidence that a Gα_s_ subunit links MIHR activation to the increase in cytoplasmic cAMP concentration in the oocyte that initiates maturation. MIHR belongs to a superfamily of neuroendocrine GPCRs involved in regulating reproduction but also nutrition, as well as diverse other physiological processes. Consistent with a wider role for *Clytia* MIH-MIHR, *MIHR* expression was detected in jellyfish tentacle cells that may respond to MIH neuropeptides produced in the gastrovascular system, while *MIHR* mutants showed sluggish behaviour and slow growth, with gamete development also stalled in jellyfish from some mutants. These findings thus open doors to an improved understanding of the vital animal process of oocyte maturation and to its evolution. More widely they shed light on the evolution of the neurohormonal regulation of sexual reproduction in the animal kingdom.

### An oocyte receptor for maturation initiation

Our work shows that *Clytia* MIHR is entirely responsible for the oocyte response to MIH, allowing us to fill in the main molecular actors leading to oocyte meiotic maturation (Fig. 5B): MIH is released from specialized gonad ectoderm cells in response to a dark-light transition at dawn via the essential photoresponsive GPCR Opsin9 [16]. Binding of MIH to MIHR at the oocyte surface is likely swiftly followed by a rise in cytoplasmic cAMP levels in the oocyte to activate cAMP-dependent Protein Kinase (PKA), as has been shown in other hydrozoan species [22]. Our antibody inhibition experiments strongly implicate Gα_s_-stimulated adenylate cyclase activity in driving this cAMP rise. PKA leads to activation of the Cdk1-CyclinB and Mos-MAP kinase systems, that have highly conserved roles in oocyte maturation [39,40]. To complete the full chain of events between MIH secretion and meiotic maturation will require the identification of PKA substrates from *Clytia* oocytes that interact with regulators of Mos translation and/or Cdk1-CyclinB autoactivation.

cAMP rises initiate oocyte maturation not only in hydrozoans but also in a diverse range of protostome and deuterostome species [6], so it is tempting to speculate that the straightforward pathway linking MIH secretion to meiotic maturation in *Clytia* may have retained its main features from a distant cnidarian-bilaterian ancestor. A useful step towards testing this idea would be to search for MIHR superfamily GPCRs in oocytes of nemertean, ascidian, bivalves or ophiuroid species that depend on cAMP signalling for maturation. If confirmed as the ancestral state, the *Clytia* MIH-MIHR system could provide a paradigm for the molecular dissection of a common, cAMP-stimulated, mechanism for oocyte meiotic resumption.

### Evolution of reproductive regulation

The role of cAMP in initiating oocyte maturation is not universal. In some species such as starfish it has no significant role [41], while in vertebrates, high cAMP levels in the ovarian oocyte maintain the prophase arrest. Clues to understanding how these marked differences arose during evolution may be found by considering the GPCRs involved in regulating gamete maturation and release across species. In vertebrates, GPCRs are involved in triggering oocyte maturation at several levels along the H-P-G axis (Figure 7). In ovarian oocytes, constitutively active GPCRs of the GPR3/6/12 family help maintain prophase arrest via Gα_s_-cAMP [12–14]. Luteinizing Hormone receptors in the surrounding follicle cells provide upstream signalling also via Gα_s_-cAMP. The follicle cells signal is transmitted to the oocyte in mouse via a cGMP decrease through gap-junctions [42], or in fish and amphibians via steroid hormone cocktails that act on one or more types of membrane receptor, potentially including GPCR progestin receptors linked to Gαi that contribute to lowering cAMP levels [39,40,43,44]. Upstream of the vertebrate follicle, LH production from the pituitary is under control of the master reproductive regulator GnRH receptor acting via Gα_q_ and cytoplasmic Ca^++^ release. Many other hormone-GPCR pairs influence production of GnRH by the hypothalamus, as well as the gonadotropins LH and FSH by the pituitary. Molecular phylogeny showed that *Clytia* MIHR is not closely to receptors of the core H-P-G GPCRs, but rather forms a superfamily with receptors of these “upstream” neuropeptide hormones, notably GnIH, QRFP, Neuropeptide Y and NkB. Production of GnRH and gonadotropins along the H-P-G axis are inhibited by GnIH and stimulated by QRFPs [45,46], while the Tachykinin family hormone NkB acts in a group of hypothalamus neurons together with Kisspeptin and Dynorphin A in the generation of cyclic GnRH pulses [47]. The ancestral cnidarian-bilaterian GPCR for MIHR and the receptors to all these vertebrate hormones may thus already had a role in regulating gamete production and/or release. In this case, vertebrate GPCRs in the hypothalamus and the pituitary, rather than those in gonad follicle cells or gametes, would have retained the trace of this ancestor. Likewise, protostome members of the MIHR GPCR group regulate sexual reproduction indirectly from sites far from the gonad. The planarian NeuropeptideY NPY-8 regulates gametogenesis through its GPCR expressed in CNS neuroendocrine hormone cells [48], while nematode Luqin peptides produced by pharyngeal neurons regulate egg laying via serotonergic RIH interneurons [49].

### An ancient regulatory system linking reproduction and nutrition?

It is striking that many bilaterian neuropeptide GPCRs closely related to *Clytia* MIHR are known for roles in regulating feeding and nutritional balance. In mammals these include GPR83/PEN [50] as well as QRFP receptors and Neuropeptide Y receptors, which provide important links between metabolic state and reproductive regulation [51–53], in conjunction with GnIH signalling [54]. Related hormone receptor families from protostomes are also known for regulating feeding, including those for *Drosophila* Leucokinin [55] and Luqins in arthropods (RYamides) and nematodes [49,56]. It is thus tempting to propose that the ancestral GPCR of the *Clytia* MIHR and these bilaterian hormone receptor families was involved in integrating sexual reproduction with nutritional status. It should be noted, however, that bilaterian neuropeptide GPCRs in this superfamily also function in many other physiological processes. Conversely, many neuropeptide hormone systems involved in integrating sexual reproduction with feeding, notably including Kisspeptin, involve GPCRs only distantly related to MIHR [35,37,57]. To unravel further the evolutionary history of hormonal regulation of gamete production in relation to nutrition it will be of great interest to examine in detail the function of MIHR in *Clytia* relating to the sluggish movement and poor growth of *MIHR* mutants, but also to determine the functions of other “group A” GPCRs from *Clytia* and other cnidarian species.

## Methods

### Animals

Sexually mature jellyfish generated from laboratory maintained *Clytia hemisphaerica* polyp colonies (“Z strains”) [26] were fed regularly with *Artemia* nauplii and cultured under light-dark cycles to allow daily spawning. Red Sea Salt brand artificial seawater (ASW) was used for all culture and experiments.

### Selection of candidate Clytia MIH Receptors

From a comprehensive *Clytia* reference transcriptome (86,606 contigs) derived from mixed larva, polyp and jellyfish samples,we generated a list of predicted protein sequences from complete and incomplete ORFs using a homemade script [58]. This dataset was screened using TMHMM 2.0c to produce a list of *Clytia* complete protein sequences coding for 7 transmembrane domain proteins (7TMD) including putative incomplete sequences containing 2 to 7 transmembrane domains. Scanning with Interproscan 5.22 was used to generate a list of 761 potential *Clytia* GPCRs with Pfam tags related to 7TMD receptors were retained, sorted by class. CD-HIT [59] was run with 95% identity to eliminate sequence duplicates, obtaining a final dataset of 536 *Clytia* GPCRs (sequences in File S1). Illlumina HiSeq 50nt reads from mRNA isolated from manually dissected gonad ectoderm, endoderm, growing oocytes and fully-grown oocytes [16] and from other *Clytia* life cycle stages including polyp and planula larvae [26] were mapped against all candidate GPCR sequences using Bowtie2 [60]. The counts for each contig were normalized per total of reads of each sample and contig length to allow expression comparisons between genes and samples.

Presumptive GPCR sequences were separated in groups using a hierarchical pipeline based on the correlation of their expressions (gplots package in R). Z-scores were obtained after standardization of the counts in the different sequenced samples for each GPCR candidate and plotted in a heat map using R (Fig. S1). For clustering, a previous collection of GPCRs[25] was complemented with several bilaterian class C GPCRs retrieved from UniProtKB and all putative *Clytia* GPCRs. Clustering analysis was performed using CLANS2 [61] with a BLOSUM62 matrix and a p-value cutoff of 1.e-30. Based on this information, Pfam signatures, reciprocal BLASTs and high expression level in oocytes, a subset of 16 top candidates was manually selected (sequences and accession numbers in FileS2).

### Candidate receptor cloning

The selected *Clytia* GPCRs were cloned from gonad-extracted cDNA into pcDNA3.1(+) (Thermo Fisher Scientific, Waltham, MA, USA) using the Gibson Assembly® Cloning Kit (New England Biolabs) [61]. pcDNA3.1(+) vector was linearized with BamHI and NotI restriction enzymes. Primers were designed using the Gibson Cloning option in Geneious v8. Forward primers consisted of the overhang left after BamHI vector linearization followed by the Kozak consensus sequence (CGCCACC), a start codon (ATG), and a sequence corresponding to the target sequence. Reverse primers consisted of the overhang left after NotI vector linearization followed by a STOP codon, and a reverse complementary sequence to the target sequence. The primers for all cloned GPCRs are listed in File S3. Polymerase chain reaction was performed using Phusion polymerase (New England Biolabs). Cloned GPCRs were sequenced using primers for T7: TAATACGACTCACTATAGGG and BGHrev: TAGAAGGCACAGTCGAGG.

### Receptor deorphanization

Cell culture GPCR ligand response assays were performed as described [28]. Briefly, CHO-K1 cells were cultured in Ham’s F12 Nut Mix medium (Thermo Fisher Scientific) with 10% foetal bovine serum. Cells were seeded in 96-well plates at approximately 10.000 cells/well and transfected the following day with pcDNA3.1(+) plasmids encoding each of the 16 candidate *Clytia* GPCRs (File S2), the promiscuous Gα-16 protein [63], and a reporter construct GFP-apoaequorin [64] (60 ng each) using the transfection reagent TurboFect (Thermo Fisher Scientific). After 2 days of expression, the medium was removed and replaced with Hank’s Balanced Salt Solution (HBSS) supplemented with 1.8 mM Ca2+, 10 mM glucose, and 1 mM coelenterazine h (Promega, Madison, WI, USA). After incubation at 37 °C for 2 hours, cells were tested by adding synthetic peptides (GenScript) in HBSS supplemented with 1.8 mM Ca2+ and 10 mM glucose. A list of all synthetic peptides used is provided in File S4. Luminescence was recorded for 45 s in a plate reader (BioTek Synergy Mx or Synergy H4; BioTek, Winooski, VT, USA). Data were integrated over the 45-s measurement period and recorded as technical triplicates in each case. Data were normalized using the response of *Platynereis* FLamide receptor to 1 μM AKYFL-NH2 [28]. Dose-response curves were obtained using concentrations between 0.01 nM and 100 μM for each peptide. Data for dose-response curves were recorded in triplicate for each concentration and the experiment was repeated independently 3 times. Dose-response curves were fitted with a four-parameter curve using Prism 6 (GraphPad, La Jolla, CA, USA) and were normalized to the calculated upper plateau values (100% activation).

### Generation of CRISPR-Cas9 mutant *Clytia* polyp colonies

Following our established protocol [65] *MIHR* small guide RNA (sgRNA) was assembled by hybridising crRNA and tracrRNA synthesized at IDT (Integrated DNA Technologies), obtaining a final concentration of 50 μM. sgRNA was kept at −80ºC until use. The crRNA *MIHR* sequence is shown in File S5. We avoided off-target matches by scanning the *Clytia* genome assembly at http://crispor.tefor.net. Purified Cas9 protein in Cas9 buffer (10 mM Hepes, 150 mM KCl) provided by J-P Concordet (MNHN Paris) was diluted to 10 μM. sgRNA was added to Cas9 protein in excess (~2:1) prior to injection and incubated for 10 minutes at room temperature. The final Cas9 concentration was adjusted to 4 μM and for sgRNA to 10 μM. The mixture was centrifuged at 14,000 rpm for 10 minutes at room temperature before injection (2-3% of egg volume) into unfertilized eggs within 1 hour after spawning, prior to fertilization.

Injected embryos were cultured for 3 days in Millipore-filtered sea water (MFSW) at 18-20°C. Metamorphosis of planula larvae into polyps was induced about 72 hours after fertilization by placing larvae (20-80/slide) on 75 × 50mm glass slides in drops of 3-4 ml MFSW containing 1 μg/ml synthetic metamorphosis peptide (GNPPGLW-amide), followed by overnight incubation. Slides with fixed primary polyps were transferred to small aquariums kept at 24°C, a temperature which favours the establishment of female colonies [66]. Primary polyps and young polyp colonies were fed twice a day with smashed *Artemia* nauplii until they were grown enough to be fed with swimming nauplii. Following colony vegetative expansion, a single well-growing colony on each slide was maintained as a founder. After several weeks of growth, polyp colonies were genotyped to assess mutation efficiency and mosaicism, and medusae were collected from the most strongly mutant colony (*MIHR* KO) for further experimentation.

### Genotyping

Genomic DNA from *Clytia* polyps was purified using DNeasy blood/tissue extraction kit (Qiagen). The *MIHR* target site was amplified by PCR using Phusion DNA polymerase. Primers used for genotyping are listed in File S5. PCR products were sequenced and mutation efficiency was assessed using TIDE analyses [64]. In cases where Agarose gel analysis of the PCR product revealed large insertions or deletions, this was cloned into pGEM easy vector and clones randomly selected for individual sequencing.

### Gonad spawning assays

Sexually mature *MIHR* mutant medusae from colonies n5-13 and n5-23 and wild type medusae were cultured on the same day-night cycle. Individual gonads were dissected in the evening after afternoon spawning. To test the light response, one group of each was transferred to 100 μl MFSW in wells of 96-well plastic plates, covered overnight and re-exposed to white light the following day. To test responses to MIH and cAMP, other groups of dissected gonads were cultured overnight in constant light, then transferred to 96-well plastic plates for two hours. MIH was added to wells as an equal volume of 200 nM WPRPamide [GenScript; 15] stock in MFSW was added to give a final concentration of 100 nM. Alternatively, a stock of 20 mM bromo adenosine 3′5′cyclic MonoPhosphate (5-Br-cAMP – Fluka; Sigma Aldrich) in distilled water was added to give a final concentration of 4 mM. Gonads were washed after 5 minutes incubation in 5-Br-cAMP. Oocyte maturation was scored after 30 minutes as Germinal Vesicle breakdown (GVBD), i.e. visible dissolution of the oocyte nuclear membrane upon entry into M phase, which in all cases was followed by spawning about 1 hour later. Gonads that showed premature maturation or spawning due to manipulation stress were excluded from analysis.

### Oocyte maturation assays

Fully grown oocytes were isolated manually from dissected gonads of wild type and *MIHR* KO mutant jellyfish prepared as above. Maturation assays were performed in wells of 96-well plates or small plastic petri dishes lined with 2% agarose in MFSW. Spontaneous maturation occurs at a low frequency in control oocytes. To test MIH-induced maturation, 10 μl WPRPamide from 1 μM solution in MFSW were added to a final concentration of 100 nM. In some experiments, 5-Br-cAMP from a 20 mM stock in H_2_O was used at a final concentration of 4 mM. GVBD was scored at 30 minutes, every 5 minutes, depending on experiments.

Antibody injection into isolated oocytes was performed using Nanoject or Narisige compressed air microinjection systems [68, 69]. Solutions were centrifuged at 14000 rpm at 4°C for 5 minutes before use, and approximately 2% oocyte volume injected. An inhibitory anti-GS antibody [33] was concentrated to 8 mg/ml in PBS by three passages through a ULTRAFREE spin column (Millipore-UFV5BQK25). A purified anti-GST antibody (Sigma 67781) at 8.3 mg/mL was used as a control.

### GPCR molecular phylogeny

Clustering analysis was performed using CLANS2 [59] with a BLOSUM62 matrix and a p-value cutoff of 1e-40 or 1e-50. Sequences were retrieved from NCBI (*Homo* and *Nematostella*) and from [28] for the deorphanized *Platynereis* GPCRs. *Nematostella* GPCRs were identified using HMMER [70]. Identifiers for the sequences used are given in Files S6 and S7. Multiple alignments were generated using Muscle [71] with default parameters. Positions containing more than 80% gaps were excluded from the alignment, provided in File S8. Phylogenetic analyses were performed using RaxML v8.2.9 [72] and the model PROTGAMMAGTR with Bootstrap support calculated from 500 replicates. The resulting tree file was visualized with FigTree (http://tree.bio.ed.ac.uk/software/figtree/).

### In situ hybridization

A urea-based protocol for *in situ* hybridization was used as previously [15,16,32].

### Immunofluorescence

For co-staining of neuropeptides and tyrosinated tubulin, dissected *Clytia* gonads, whole medusae were fixed overnight at 18ºC in HEM buffer (0.1 M HEPES pH 6.9, 50 mM EGTA, 10 mM MgSO_4_) containing 3.7% formaldehyde, then washed five times in PBS containing 0.1% Tween20 (PBS-T). Treatment on ice with 50% methanol/PBS-T then 100% methanol plus storage in methanol at −20ºC improved visualization of microtubules in neural cells. Samples were rehydrated, washed several times in PBS-0.02% Triton X-100, then one time in PBS-0.2% Triton X-100 for 20 minutes, and again several times in PBS-0.02% Triton X-100. After overnight incubation at 4ºC in PBS with 3% BSA they were incubated in a rabbit anti-PRPa antibody [15] and a rat monoclonal anti-Tyr tubulin (YL1/2, Thermo Fisher Scientific) in PBS/BSA at room temperature for 2 h. After washes, the specimens were incubated with secondary antibodies (Rhodamine goat anti-rabbit and Cy5 donkey anti-rat-IgG; Jackson ImmunoResearch, West Grove, PA) overnight in PBS at 4ºC, and nuclei stained using Hoechst dye 33258. Images were acquired using a Leica SP8 confocal microscope and maximum intensity projections of z-stacks prepared using ImageJ software.

### Statistics

Fisher exact tests were performed at http://www.socscistatistics.com.

## Non-standard Abbreviations

*MIH*: Maturation Inducing Hormone
*GVBD*: Germinal Vesicle Breakdown

## Acknowledgements

We thank our LBDV colleagues Philippe Dru for Bioinformatics support, Maeva Goulais for assistance with immunofluorescence and Céline Hebras for antibody injections. Thanks also to J-P Concordet (MNHN Paris) for generously providing Cas9 protein, Noriyo Takeda (Hiroshima University) for the anti-PRPamide antibody and Laurinda Jaffe (University of Connecticut Health Centre) for the anti-Gα_S_ antibody. Funding was provided by the Marie Curie ITN NEPTUNE and French ANR grant OOCAMP–ANR-13-BSV2-0008, as well as core CNRS and Sorbonne University funding to the LBDV. For animal care and microscopy we thank of the Service Aquariologie (especially Alexandre Jan) and the Villefranche-sur-mer Imaging platform of the Institut de la Mer de Villefranche (IMEV)/EMBRC-France (ANR-10-INBS-02).

## List of Supplementary Figures and Files

Figure S1. Expression of *Clytia* GPCRs across tissues and life cycle stages.

Figure S2. Phylogenetic tree file corresponding to Figure 6D

File S1. FASTA file of *Clytia* candidate GPCR sequences

File S2. FASTA file of cloned *Clytia* candidate MIHR sequences File S3. PCR primers for GPCR cloning

File S4. Peptides used for GPCR deorphanisation

File S5. crRNA CRISPR and PCR primers for mutant genotyping File S6. Sequences used for CLANS2 clustering in Figure 6A and B

File S7. Sequences used for CLANS2 clustering in Figure 6C

File S8. FASTA file of the GPCR alignment used for phylogenetic analyses shown in Figure 6D

