## Supplementary Figures for "A G-protein-coupled receptor mediates neuropeptide-induced oocyte maturation in the jellyfish *Clytia*"

**Figure S1. Expression of *Clytia* GPCRs across tissues and life cycle stages.**

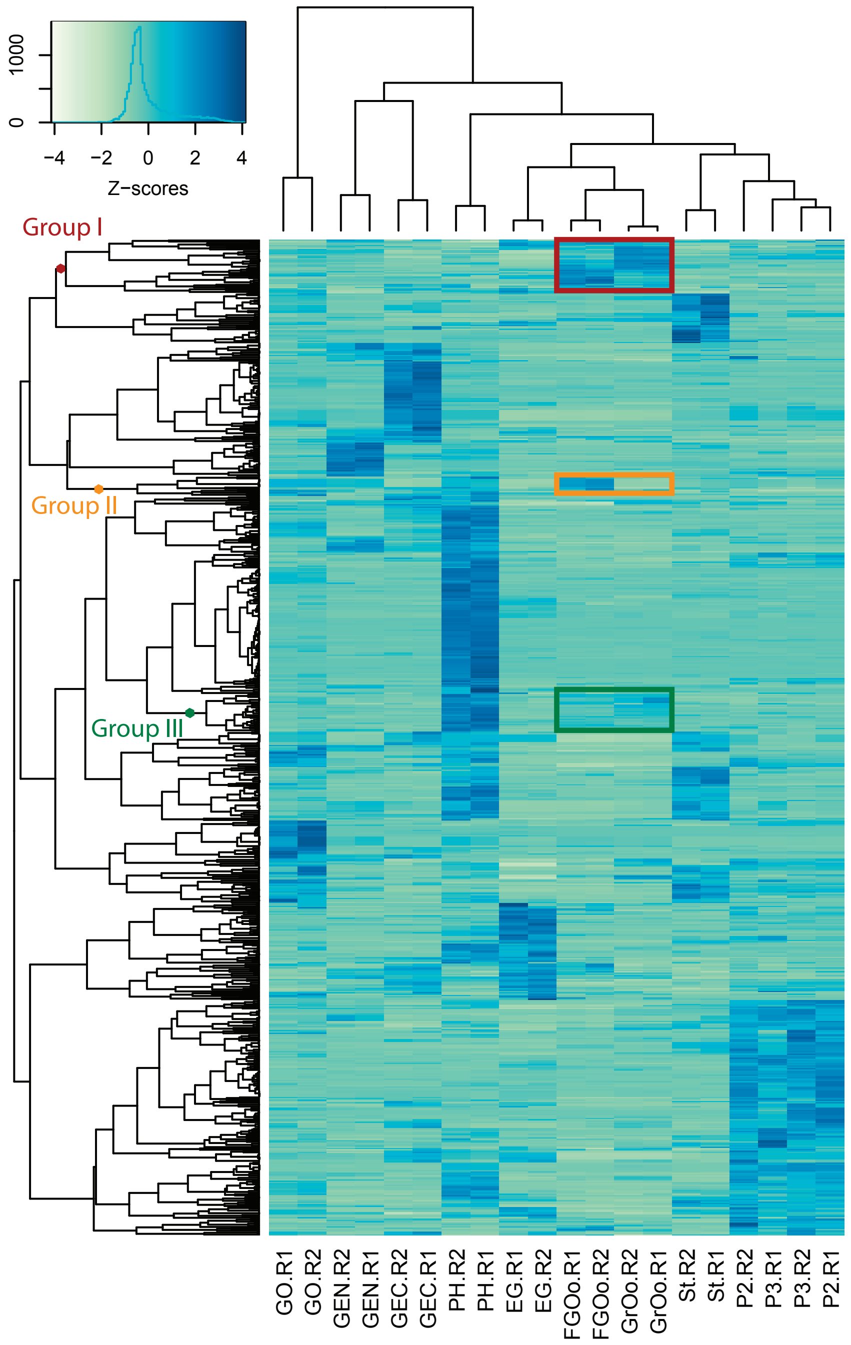


Heat map representing the expression of putative *Clytia* GPCRs in the different samples studied derived from RNA-seq data (see methods), in which sequences are clustered according to similarity of their profiles across tissues and stages. Z-score values are colour-coded to reflect significantly higher (dark blue) or lower (pale green) than average values -see z value distribution in inset. Three main profile groups showed expression enriched in the oocytes (coloured boxes). Abbreviations: R1/R2= Biological Replicate1/2 for Illumina sequencing; GO= Gonozooid; GEN= Gonad Endoderm; GEC= Gonad Ectoderm; PH= Polyp Head; EG= Early Gastrula; FGOo= Fully Grown Oocytes; GrOo= Growing Oocytes; St= Stolon; P2/3= 2/3-day old planula larvae.

**Figure S2. Phylogenetic tree file corresponding to Figure 6D**

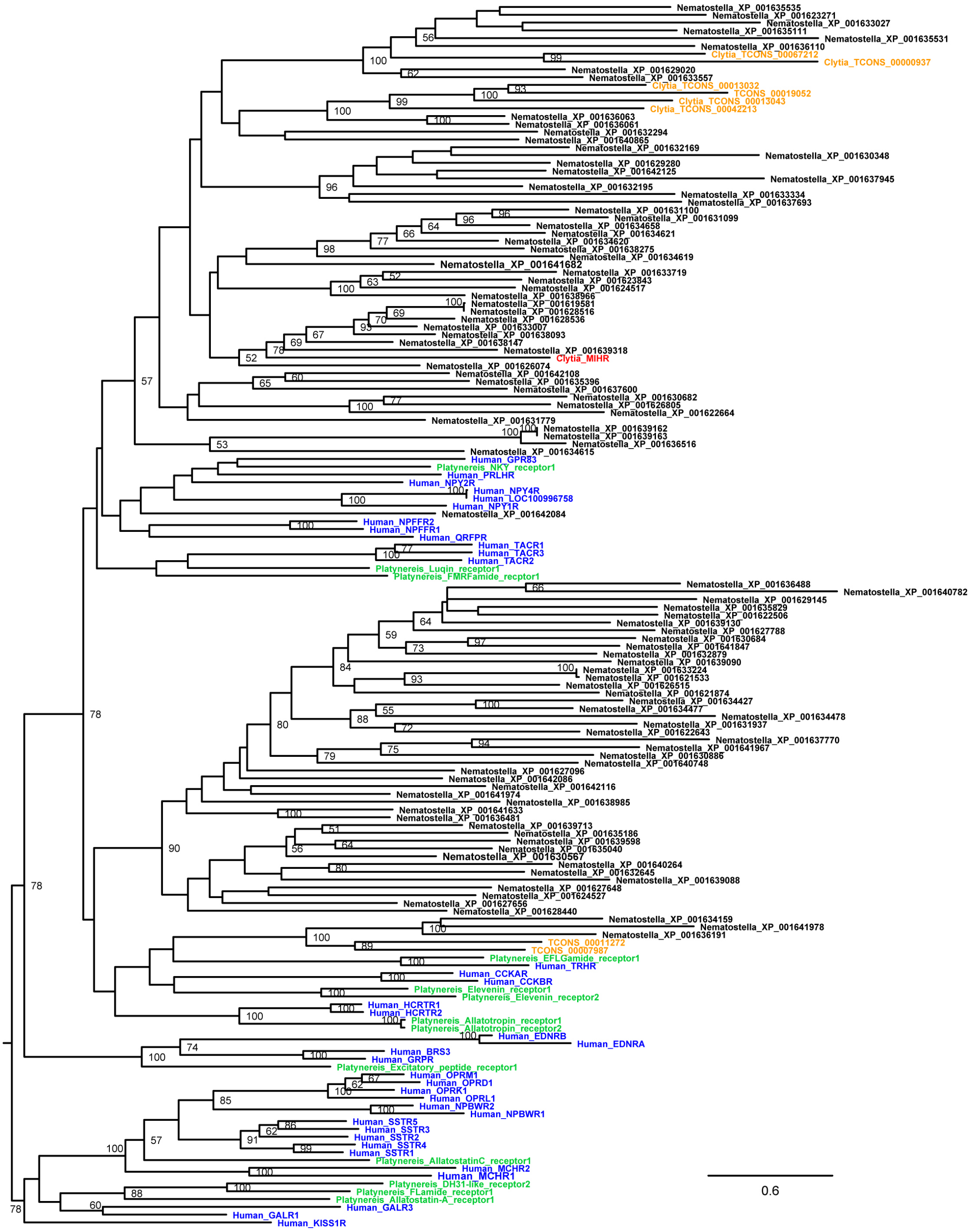


Heat map representing the expression of putative *Clytia* GPCRs in the different samples studied derived from RNA-seq data (see methods), in which sequences are clustered according to similarity of their profiles across tissues and stages. Z-score values are colour-coded to reflect significantly higher (dark blue) or lower (pale green) than average values -see z value distribution in inset. Three main profile groups showed expression enriched in the oocytes (coloured boxes). Abbreviations: R1/R2= Biological Replicate1/2 for Illumina sequencing; GO= Gonozooid; GEN= Gonad Endoderm; GEC= Gonad Ectoderm; PH= Polyp Head; EG= Early Gastrula; FGOo= Fully Grown Oocytes; GrOo= Growing Oocytes; St= Stolon; P2/3= 2/3-day old planula larvae.
